# Improving Inverse Folding models at Protein Stability Prediction without additional Training or Data

**DOI:** 10.1101/2024.06.15.599145

**Authors:** Oliver Dutton, Sandro Bottaro, Michele Invernizzi, Istvan Redl, Albert Chung, Falk Hoffmann, Louie Henderson, Stefano Ruschetta, Fabio Airoldi, Benjamin M J Owens, Patrik Foerch, Carlo Fisicaro, Kamil Tamiola

## Abstract

Deep learning protein sequence models have shown outstanding performance at *de novo* protein design and variant effect prediction. We substantially improve performance without further training or use of additional experimental data by introducing a second term derived from the models themselves which align outputs for the task of stability prediction. On a task to predict variants which increase protein stability the absolute success probabilities of ProteinMPNN and ESMif are improved by 11% and 5% respectively. We term these models ProteinMPNN-ddG and ESMif-ddG.

## 1 Introduction

Models trained to predict native protein sequence based off of structural and partial sequence context show remarkable versatility. They generalise both to the macro scale, designing whole proteins, and the micro scale, predicting beneficial single point mutations without explicit training [1, 2]. Previous work has improved performance of ProteinMPNN [3] at designing soluble analogues of membrane proteins by retraining with a modified training set, while AlphaMissense [4] improved the accuracy of AlphaFold2 [5] at distinguishing disease related mutations by fine-tuning with unlabelled data. The present work continues in this direction but without the usage of retraining or fine-tuning, instead predictions are made with two different inputs and the difference in outputs are utilised.

Unsupervised deep learning models leveraging sequence and/or structural information have demonstrated zero-shot prediction of the change in protein properties upon mutation, including expression, activity and stability [2, 6, 7, 8, 9]. Models leveraging only sequence data have been shown to outperform those incorporating structural data on the majority of properties except for stability [8]. The effect of point mutations on stability was best predicted by inverse folding models which are trained to recover the native sequence of a protein using its backbone structure and partial sequence context. The differences between the likelihood of the native and a mutant amino acid predicted by these models correlate to the relative stability of the two sequences.

By considering toy cases, we find that exact predictions from an inverse folding model are non-optimal for the prediction of physical quantities such as protein stability. We introduce an additional term derived from the model itself to improve the prediction of relative stability upon mutation without experimental data or further training. We demonstrate this modification improves accuracy in predicting the effect of point mutations for two popular models, ProteinMPNN and ESMif [1, 2], across three benchmark datasets. We term the resulting improved, unsupervised models ProteinMPNN-ddG and ESMif-ddG.

In addition, we implement a novel tied decoding scheme which limits the increased computation cost of ProteinMPNN-ddG relative to common usage of ProteinMPNN.

## 2 Specialisation of model for mutation stability prediction

### 2.1 Leveraging maximum sequence context

We aim to predict the effect of a mutation from the wild type amino acid, *X*, to a mutant, *Y* , at position *i*, denoted *f*_*i*,*X*_→_*Y*_ , using a subset of the sequence, *s*, and positions of the backbone atoms, *x*. Following common usage of large language models [7, 10, 2] we mask the identity of the amino acid at position *i* and predict using all other possible sequence and structural context (Equation 1). In the baseline models we score mutations as

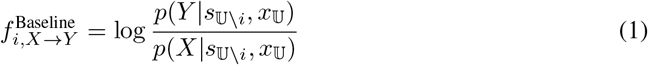

where 𝕌 denotes the set of all residues and 𝕌 \ *i* is the set of all residues with residue *i* masked. The log-odds ratio are calculated using either ProteinMPNN or ESMif.

In its naïve usage, ProteinMPNN only uses a subset of the full sequence context for predicting most tokens as residue identities are decoded autoregressively with a random order, hence the first residue decoded has no sequence context while on average only half the sequence context is given. We define this as our baseline ProteinMPNN in the results described. Here, we generate separate decoding orders for each residue such that it is decoded last, so is predicted with full sequence context, and find this procedure improves accuracy over the classic usage, improving sequence recovery on the ProteinMPNN validation set by nearly 4% (Table 1). A single structure for each of the 1,464 clusters in the ProteinMPNN validation set was taken and predicted, resulting in predictions for 375,445 residues on which sequence recovery metrics were computed in Table 1. As the predictor ESMif is autoregressive from N to C terminus, only sequence context from residues before position *i* can be given so we use *s* {_0,1,…,*i* − 1_} in place of *s*_𝕌 /*i*_ for ESMif. With retraining this limitation could be removed, but we do not pursue it in this work.

**Table 1:**
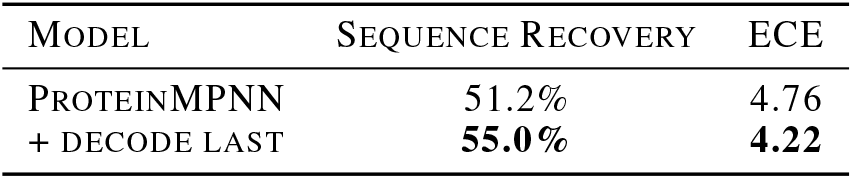
Improved sequence recovery metrics from tailored usage of ProteinMPNN with Exponen-tiated mean Cross-Entropy (ECE) computed over 20 natural amino acids.

| MODEL | SEQUENCE RECOVERY | ECE |
| --- | --- | --- |
| PROTEINMPNN | 51.2% | 4.76 |
| + DECODE LAST | **55.0%** | **4.22** |

### 2.2 Utilising logit perturbation between different inputs

In the limiting case where no structural or sequence context are given, an optimal model would return a distribution mirroring the amino acid frequencies in its training set. This would lead to log-odds ratios suggesting mutations to more abundant amino acids, such as Tryptophan to Leucine, are more likely to increase stability than Leucine to Tryptophan. However, the natural abundance of amino acids in proteins has been shown to be linked to a trade-off between the metabolic costs associated with each amino acid and the complexity of the proteome that composition generates [11], rather than to an intrinsic ability to stabilise a protein structure. Motivated by this observation, we shift the log-odds ratios such that mutations are predicted to have no effect when no context is given. This decision is not based on any experimental stability data.

Furthermore, if only the backbone atoms of a single residue are given, a model can also utilise discrepancies in the relative geometry of the backbone atoms. This backbone residue internal geometry gives information about the amino acid identity but not stability, as protein stability arises from the interactions between amino acids. Hence, we recommend corrections to nullify this contribution to the models predictions.

We therefore introduce an additional term consisting of the log-odds ratio as in Equation (1) but where only backbone atoms of the single residue being predicted are given as context:

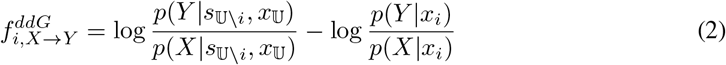

We note Equation (2) resembles the standard procedure in classical free energy calculations in which the free energy change upon mutation is estimated both in folded and unfolded states [12]. In this analogy the unfolded state, the second term, is modelled as the singular amino acid.

We first verified that ProteinMPNN and ESMif generalise to single residue inputs. We calculated averaged predictions for each of the 380 possible *X* → *Y* mutations as *δ*_*X*→*Y*_ :

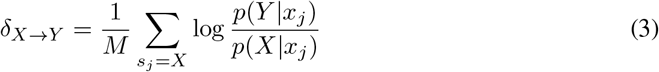

where the sum runs over *M* structures in a structural database. A single structure was taken for each of the 23,349 clusters in the training set of ProteinMPNN, and residues for which the backbone atom positions are unknown were removed. This resulted in 5,615,050 residue geometries with at least 77,000 geometries for each distinct amino acid. The values for ProteinMPNN are shown in Figure A.1a. We found the log-odds ratios calculated from the amino acid frequencies in the training set of ProteinMPNN correlate well with averaged log-odds ratios 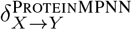 (spearman correlation coefficient 0.72).

As log-odds ratios calculated from the amino acid frequencies are naturally antisymmetric, the only possible origin of deviation from antisymmetry, |*δ*_*X*→*Y*_ +*δ*_*Y*→*X*_| , is structural context derived from the backbone atoms N, C*α*, C and O geometry. As expected, the largest size of deviation from antisymmetry for ProteinMPNN were observed for Glycine, Proline, Valine and Isoleucine (Figure A.1b). Looking at the backbone N-C*α*-C opening angle, we observe most amino acids display a very similar distribution, while Glycine, Proline, Valine and Isoleucine are quite distinct (Figure A.2). Glycine due to its lack of sidechain, and Proline due to its ring incorporating C*α* and N, display increased N-C*α*-C opening angle. Valine and Isoleucine are the only amino acids with multiple alkyl groups attached to C*β*, which causes steric crowding around C*α*, leading to a smaller N-C*α*-C opening angle. The clear relationship between amino acid frequencies in the training set and log-odds ratios in *δ*_*X*→*Y*_ in addition to pronounced deviation from antisymmetry, |*δ*_*X*→*Y*_ + *δ*_*Y*→*X*_| , for amino acids with distinct geometries demonstrates that ProteinMPNN generalises well to single residue inputs. We performed a similar analysis for ESMif and, after introducing a small correction due to the autoregressive nature of the model (see Appendix A.1), we find analogous results.

### 2.3 Time complexity reduction to enable proteome-scale prediction with ProteinMPNN-ddG

ProteinMPNN implements a causal decoding scheme of protein sequence identity using any order. This means the first residue to be decoded has purely structural context with no sequence information while the final one to be decoded is given the sequence information on all other residues. By running the model separately for each residue with an appropriate decoding order we can predict sequence probabilities for every residue with full context, increasing native sequence recovery (Table 1).

Naïvely decoding every residue with full context requires *N* passes of ProteinMPNN, where *N* is the number of residues in the protein. We analysed the model to find any shared work between passes. The model involves three stages:

1. Nearest neighbors computation for each residue, these define edges in the sparse graph
2. Encoder of structural features
3. Decoder using both sequence and structural features

The decoding order of sequence information is only relevant for the final stage, so the first two stages only need to be computed once and reused with different decoding orders. We implemented a version of ProteinMPNN with shared work and find this displays linear slowdown of approximately *N/*5 relative to a single pass rather than the full *N* fold on an NVIDIA A100 40GB GPU. However, this improvement alone is not enough to allow cheap mutation stability prediction at scale.

We made further improvements inspecting our aim for full sequence context in Equation (1). The only requirement on the decoding order for predicting residue *i* is that it is decoded last, there are no constraints placed on the order up until then. Acceptable orders to predict residues 0 and 1 can share the same order up until the last two, e.g. in an 8 residue system the orders 23456710 and 23456701 can be used with the partial decoding of 234567 shared. This principle was extended to maximise the amount of shared partial decoding, which reduces the number of token decodes from 𝒪 (*N* ^2^) when random orders are used to 𝒪 (*N* log *N* ) (Figures A.3 and A.4). This tied decoding greatly reduced the additional cost associated with using full sequence context relative to a single pass of ProteinMPNN from over 200-fold for a 1024 residue protein, to under 4-fold (Figure A.5). This speedup enables identification of mutations affecting protein stability at the proteome-scale. Saturation mutagenesis predictions were made for all 23,391 AlphaFold2 predicted structures of the human proteome (UP000005640_9606_HUMAN) in 30 minutes on a single V100 16GB GPU.

ProteinMPNN-ddG represents a million-fold speedup relative to established predictors such as FoldX and Rosetta [13, 14] on similar hardware, and a four-hundred-fold speedup relative to Rapid Stability Prediction (RaSP) [15], which is trained to approximate Rosetta predictions at a lower computational cost (Table 2). Throughput for ProteinMPNN-ddG was computed as an average over processing the whole proteome (including compilation and pdb loading times) on a single NVIDIA V100 16 GB GPU machine. Throughputs for RaSP, Rosetta and FoldX taken from highest throughput results in [15], where RaSP runtime is calculated on single NVIDIA V100 16 GB GPU machine while Rosetta and FoldX computations were parallelized and run on a server using 64 2.6 GHz AMD Opteron 6380 CPU cores with three ΔΔ*G* computations per mutation.

**Table 2:** Throughput of ProteinMPNN-ddG and other methods.

| MODEL | THROUGHPUT (RESIDUES/SECOND) |
| --- | --- |
| PROTEINMPNN-DDG (OURS) | **9800** |
| RASP | 2.5 |
| ROSETTA | 0.0052 |
| FOLDX | 0.0025 |

## 3 Experiments

### 3.1 Datasets

We consider three protein stability datasets used in previous studies:

1. Tsuboyama *et al*. [16] produced a saturation mutagenesis dataset measuring resistance to proteolytic digest, which has been shown to correlate well with thermal stability. We follow the subset selected in [15], where only single amino acid substitutions with high quality ΔΔ*G* values for natural proteins were used, comprising of 164,524 point mutations across 164 proteins.
2. S2648 dataset contains 2,648 point mutations spread across 131 proteins, with values curated from the ProTherm database [17]. S2648 has been used to train ΔΔ*G* predictors in previous studies [18, 19, 20].
3. S669 [21] contains 669 variants of 94 proteins with less than 25% sequence identity to those in S2648 such that it can function as a test dataset for models trained on S2648.

### 3.2 Metrics

A common task in protein engineering is to identify protein variants with increased thermodynamic stability. We therefore focus on differentiating sequences with increased stability (ΔΔ*G <* 0), mirroring real world usage as closely as possible, subject to constraints imposed by the size and sparsity of available datasets.

Often only a manageable handful (∼10) single mutations are experimentally measured which corresponds to less than 1% of the possible options for a *>*50 residue protein. This selection can be made e.g. by identifying the top-scoring mutations from all possible options. We define a metric, ‘Success@10’, where the 10 single-point mutations with highest predictions for each protein are selected and the proportion which displays higher stability than the wild type protein is calculated. The ‘Success@10’ metric is meant to reflect practical usage of such models. As the Tsuboyama dataset contains data for nearly all possible point mutations for 164 proteins, the Success@10 metric is averaged over 1,640 data points so is statistically meaningful. In line with previous works, for S2648 and S669 datasets where a low number of data points are available per protein we compute the area under the receiver operating characteristic curve (auROC) and area under precision recall curve (auPRC) for differentating mutations which experimentally show increased stability relative to the wild type protein. Note that auROC and auPRC do not take into account that predictions at the top end of the score distribution are more relevant to real world usage. Metrics such as Normalized Discounted Cumulative Gains, NDCG [8, 22, 23] have been proposed to increase the weighting of high scoring mutations, however the selection of discount rate in NDCG is required.

Thanks to the large number of mutations per protein approximating saturation mutagenesis in the Tsuboyama dataset, we can compute statistically meaningful metrics on a per-protein basis then average over all proteins, as is common practice for deep mutation scanning datasets [24, 8]. Due to the low number of mutations for each protein in S2648 and S699, metrics must be computed over the aggregate of all wild type proteins present.

### 3.3 Benchmark models

We compared the performance of ProteinMPNN-ddG and ESMif-ddG with publicly available methods that enable large-scale predictions at an affordable computational cost. Since the focus of this work is on improving unsupervised inverse folding models at mutation prediction tasks, we only consider few supervised models (RaSP [15], DDGun [25], ACDC-NN [26]). Details about the chosen benchmark models are provided in the appendix A.3.

## 4 Results

Both ProteinMPNN-ddG and ESMif-ddG showed improvements across all datasets and metrics, Table 3. In the challenging task of selecting the top 10 single-point mutations for each protein and measuring the proportion that exhibit higher stability than the wild-type protein (‘Success@10’), our modifications increased absolute success rates by 11% (from 66% to 77%) for ProteinMPNN-ddG and by 5% (from 64% to 69%) for ESMif-ddG. This is a significant improvement for real-world applications, achieved without additional model training or data, while maintaining a compute efficiency of up to 10,000 residues per second on a NVIDIA V100 16 GB GPU.

**Table 3:** Accuracy of predictions for various models and datasets.

| MODEL | TSUBOYAMA, S164524 |  |  | S2648 |  | S669 |  |
| --- | --- | --- | --- | --- | --- | --- | --- |
|  | SUCCESS@10 | AUROC | AUPRC | AUROC | AUPRC | AUROC | AUPRC |
| *Unsupervised* |  |  |  |  |  |  |  |
| PROTEINMPNN | 66% | 0.78 | 0.49 | 0.75 | 0.44 | 0.64 | 0.36 |
| PROTEINMPNN-DDG (OURS) | **77%** | **0.82** | **0.56** | **0.80** | **0.52** | **0.71** | **0.44** |
| ESMIF | 64% | 0.77 | 0.47 | 0.69 | 0.41 | 0.63 | 0.39 |
| ESMIF-DDG (OURS) | 69% | 0.79 | 0.50 | 0.72 | 0.45 | 0.67 | 0.41 |
| *Supervised* |  |  |  |  |  |  |  |
| RASP | 51% | 0.73 | 0.41 | 0.76 | 0.43 | 0.70 | 0.43 |
| ACDC-NN-SEQ | 51% | 0.70 | 0.38 | 0.75 | 0.43 | **0.71** | 0.41 |
| ACDC-NN-STRUCT | 42% | 0.71 | 0.38 | 0.76 | 0.40 | **0.71** | **0.44** |
| DDGUN | 50% | 0.69 | 0.37 | 0.70 | 0.39 | 0.69 | 0.36 |
| DDGUN3D | 41% | 0.70 | 0.36 | 0.73 | 0.38 | 0.70 | 0.40 |

Ablations on ProteinMPNN show improvements from both the usage of full sequence context and nullifying background effects by subtracting predictions derived from single residue context (Table A.2). We also demonstrate that using predictions from single residue context outperforms simply shifting by the amino acid frequencies in the ProteinMPNN training set.

The general approach presented here, which consists of leveraging difference in model outputs from different inputs, can be extended beyond stability prediction. We note that the change in logit differences between predictions for a protein with and without its binding partner can accurately differentiate mutations which strengthen binding. We leave a thorough benchmark of this to later works.

## A Appendix

### A.1 Correction to ESMif-ddG

When single residue inputs are given to ESMif, it infers them as the first residue in the sequence. This leads ESMif to more strongly reflect amino acid frequencies of the first position in its training set rather than overall aboundances. Over 98.5% of the training set is made up of AlphaFold2 struc-tures predicted for UniRef50 sequences [2]. These UniRef50 sequences all begin with methionine, being coded at the mRNA level by the canonical eukaryotic start codon AUG. As a result, we observe predictions with a single residue context show high probabilities of methionine. ProteinMPNN does not use absolute positional information and does not appear to suffer from this behaviour. We observed that the non-methionine *δ*_*X*_→_*Y*_ for ESMif correlated well to those from ProteinMPNN (pearson correlation coefficient 0.83), and we therefore utilised ProteinMPNN predictions to derive a correction for the methionine bias of ESMif. We fitted one coefficient, representing the background over-prediction of ESMif for methionine, shifting the related parameters for *M*→*X* and *X*→*M* by adding or subtracting the coefficient and maximising the pearson correlation coefficient between the ProteinMPNN and ESMif *δ*_*X*→*Y*_ . The pearson correlation coefficient between 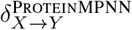 and 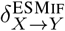 *δ*_*X*→*Y*_ improved from 0.52 to 0.85 after training the single methionine coefficient, which affects only 38 of the 380 values in *δ*_*X*→*Y*_ . We applied this correction to all ESMif predictions, following the equation

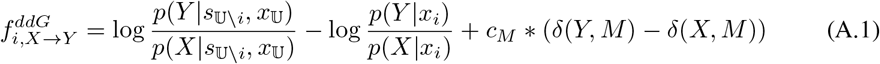

where *M* is methionine, *c*_*M*_ = 4.18 is the trained coefficient and *δ* is the Dirac delta function.

For completeness, we report results excluding mutations involving methionine (Table A.1) in addition to the complete ones of Table 3.

**Table A.1:** Results with all mutations involving methionine removed.

| MODEL | TSUBOYAMA, S164524 |  |  | S2648 |  | S669 |  |
| --- | --- | --- | --- | --- | --- | --- | --- |
|  | SUCCESS@10 | AUROC | AUPRC | AUROC | AUPRC | AUROC | AUPRC |
| *Unsupervised* |  |  |  |  |  |  |  |
| PROTEINMPNN | 65% | 0.78 | 0.50 | 0.75 | 0.44 | 0.65 | 0.35 |
| PROTEINMPNN-DDG (OURS) | **76%** | **0.82** | **0.56** | **0.80** | **0.52** | **0.71** | 0.42 |
| ESMIF | 64% | 0.77 | 0.47 | 0.70 | 0.41 | 0.64 | 0.39 |
| ESMIF-DDG (OURS) | 69% | 0.79 | 0.50 | 0.72 | 0.45 | 0.68 | 0.40 |
| *Supervised* |  |  |  |  |  |  |  |
| RASP | 51% | 0.73 | 0.41 | 0.76 | 0.42 | 0.69 | 0.42 |
| ACDC-NN-SEQ | 49% | 0.70 | 0.37 | 0.75 | 0.43 | 0.70 | 0.39 |
| ACDC-NN-STRUCT | 42% | 0.71 | 0.37 | 0.76 | 0.39 | 0.70 | **0.43** |
| DDGUN | 49% | 0.69 | 0.37 | 0.71 | 0.39 | 0.69 | 0.34 |
| DDGUN3D | 41% | 0.70 | 0.35 | 0.73 | 0.38 | 0.69 | 0.38 |

### A.2 Approximate theoretical slowdown of decode last

We estimated the relative cost of the tied decoding scheme to a single pass of ProteinMPNN. ProteinMPNN is a message passing neural network on a sparse graph, where each node has at most 48 edges. The bound number of edges per node leads to constant decode cost per residue. The encoder and decoder stages share the same hidden dimension and number of layers, though the decoder updates both edges and nodes while the encoder only updates nodes. We approximate decoder and encoder as equal cost. The encoder stage is independent of decoding order so has no increased cost, with *N* decodings for both. In the decoder stage the number of decodings increases from *N* to (*N* log_2_ *N* + *N* ) in the tied decoding scheme if *N* is a power of two. This gives a theoretical bound of the slowdown to use full sequence context rather than partial as 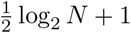 (Equation A.2). Our timings benchmark were all within theoretical bounds (Figure A.5).

The relative slowdown of tied decoding to a single pass, S, is given by:

**Figure A.1:**
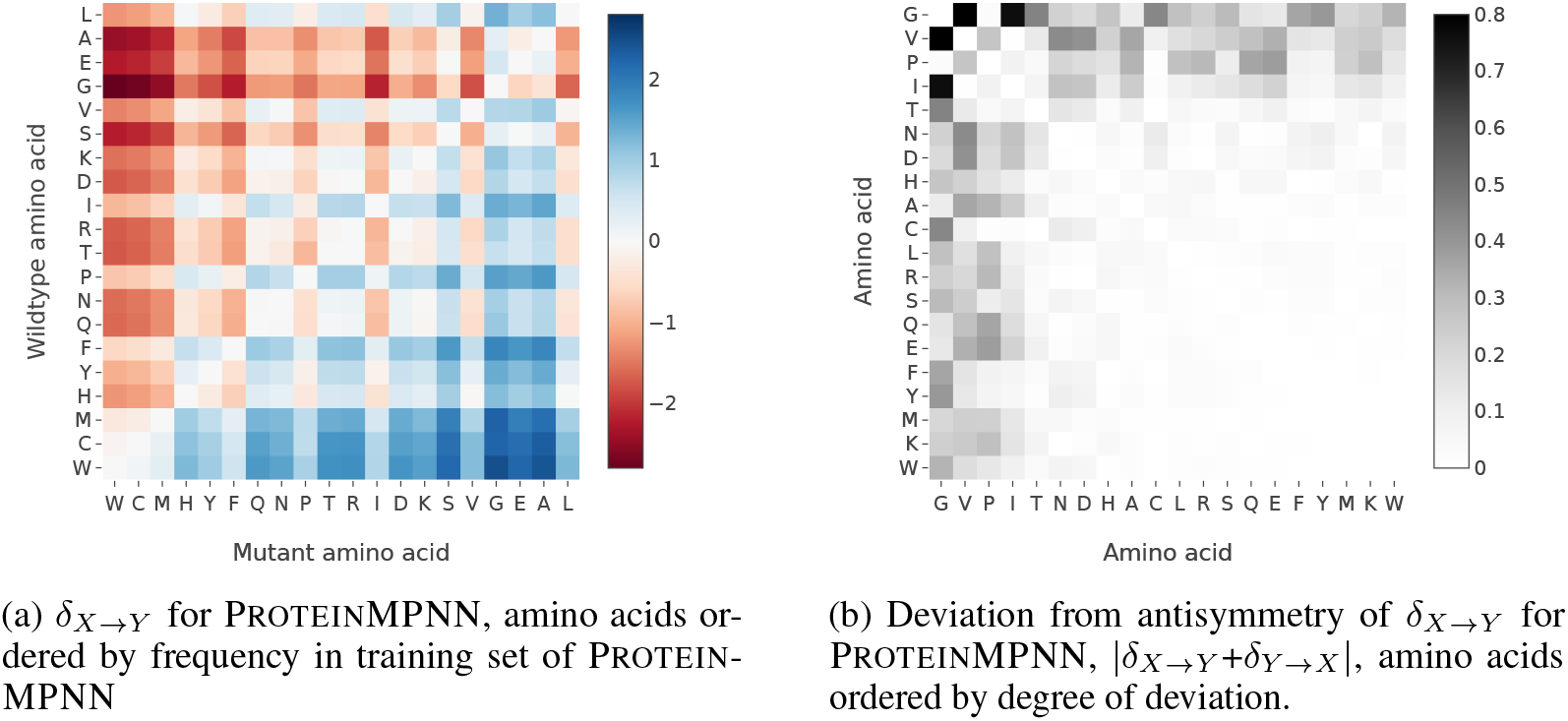
Values and deviation from antisymmetry for *δ*_*X*→*Y*_ of ProteinMPNN

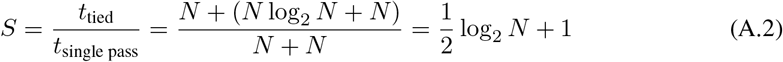

**Figure A.2:**
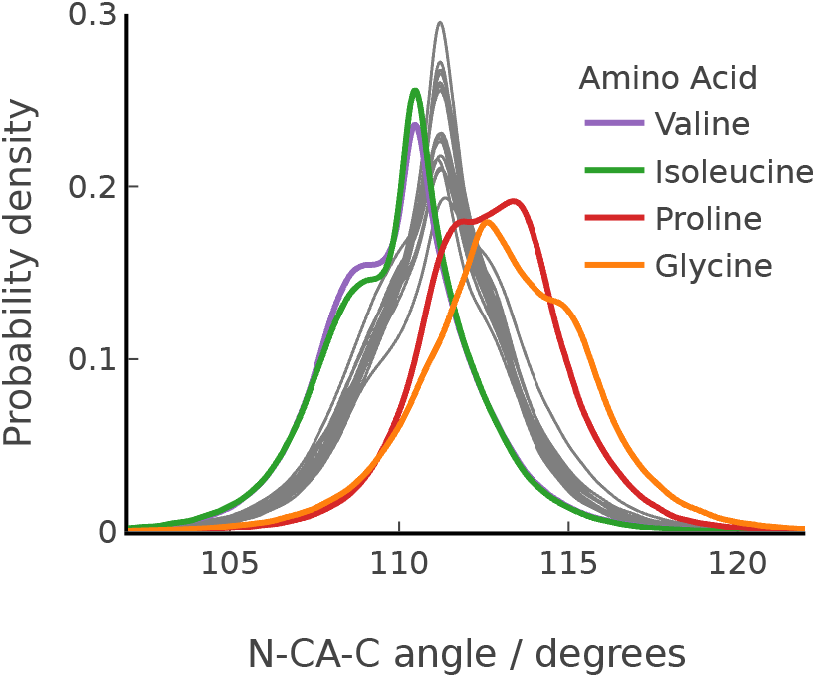
N-C*α*-C angle distributions for all 20 natural amino acids

### A.3 Benchmark models

As this work is focused on methods to improve unsupervised inverse folding models at mutation effect predictions tasks, in particular stability, we only benchmark a small subset of supervised models available in literature. The chosen models are all predictors with public code and strong results in literature, which could be scaled to predict *>* 100, 000 point mutations within a reasonable computational budget. We did not consider here the commonly used ΔΔ*G* predictors Rosetta and FoldX as they both did not display state-of-the-art performance in previous benchmarks and due to high computational cost [15].

Here are the models considered for our benchmark, reported in Table 3.

- Rapid Stability Prediction (RaSP) [15] is a deep learning predictor trained to reproduce the predictions of Rosetta, a well-established energy-function based method [27, 28].
- DDGun [25] is a mutant effect predictor based off of a linear combination of features shown to correlate to protein stability, weighted by their correlation on a mutant stability dataset. Here we consider both the sequence based model, as well as the structure and sequence based model DDGun3D. It leverages the multiple sequence alignment (MSA) for the protein being predicted which we generated using hhblits [29] based on the June 2020 edition of UniRef30 where tabulated predictions were not found.
- ACDC-NN [26] is a deep neural network trained on S2648 demonstrating strong performance on a related dataset, outperforming energy-based methods such as Rosetta and FoldX as well as DDGun [13]. We here consider both the sequence based model (ACDC-NN-Seq), as well as the structure and sequence based model ACDC-NN-Struct. This model leverages an MSA which was generated according to instructions on the associated GitHub page^1^, if tabulated predictions were not found.

**Figure A.3:**
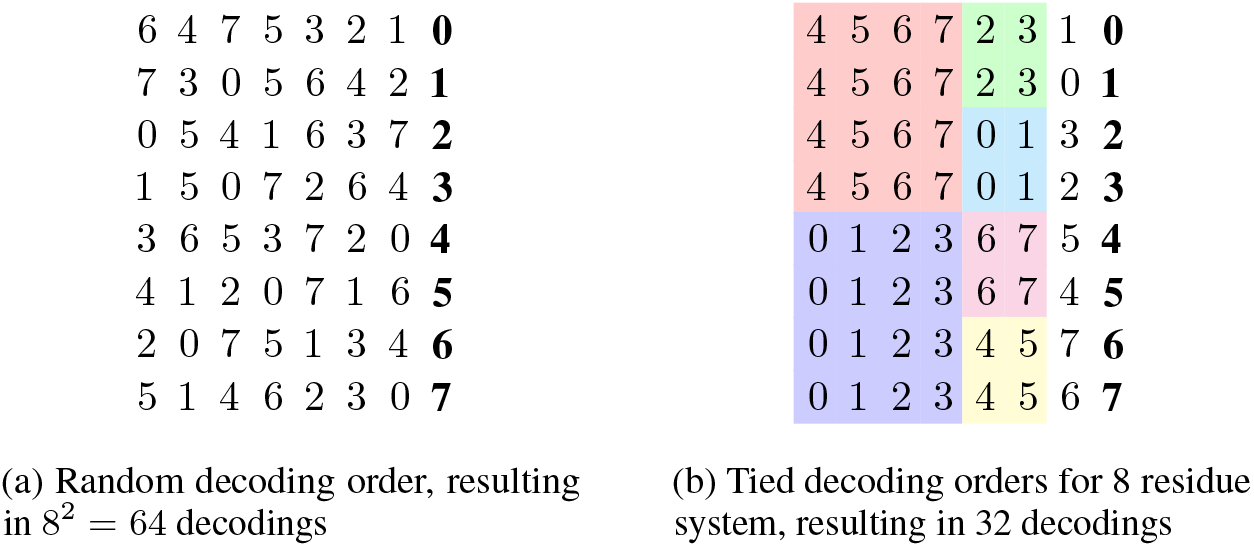
Comparison of decoding schemes which end in a particular residue being decoded last which are random vs tied to minimise compute required, decoding proceeds from left to right, with different final residues decoded in the far right column

**Figure A.4:**
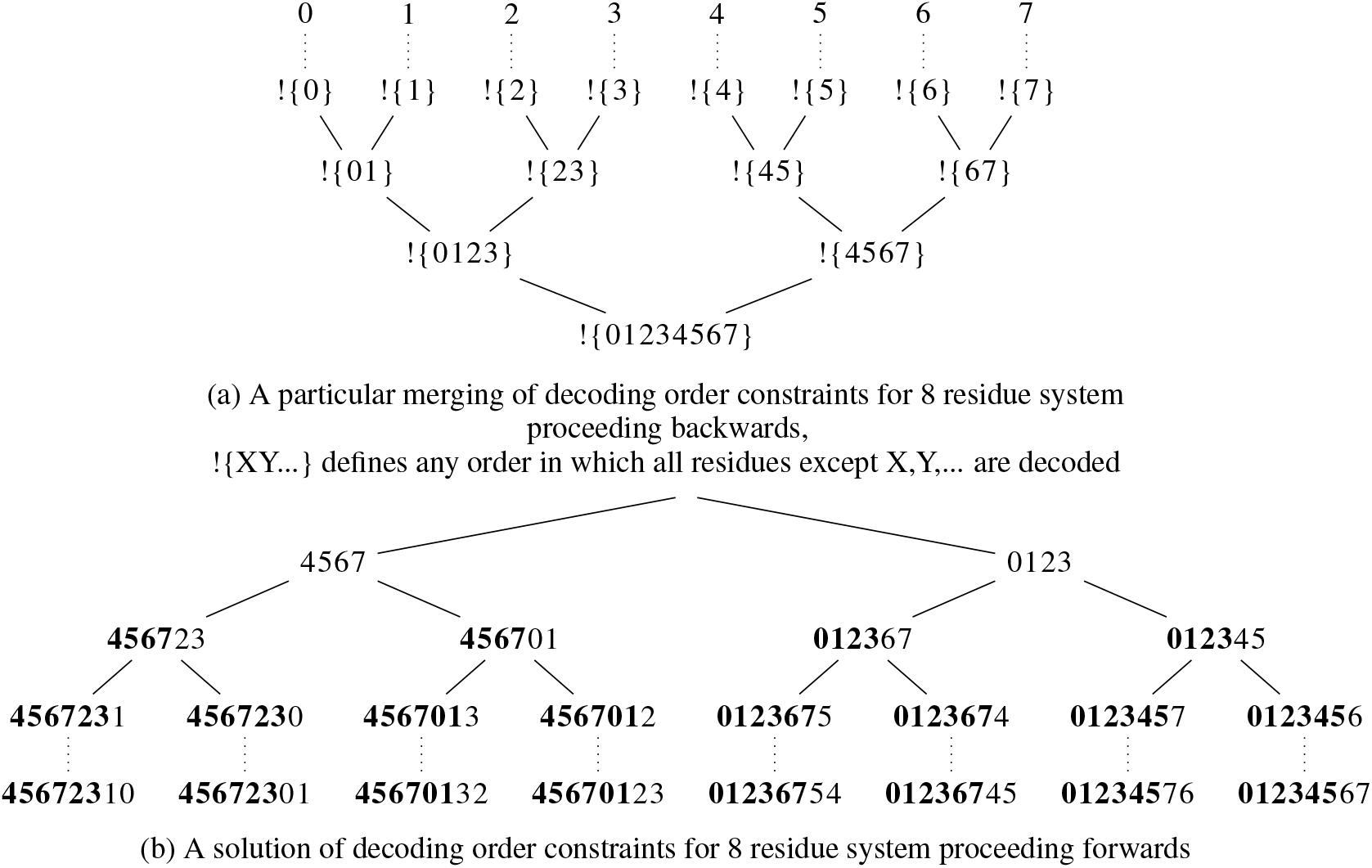
Shared decoding order visualisation

**Figure A.5:**
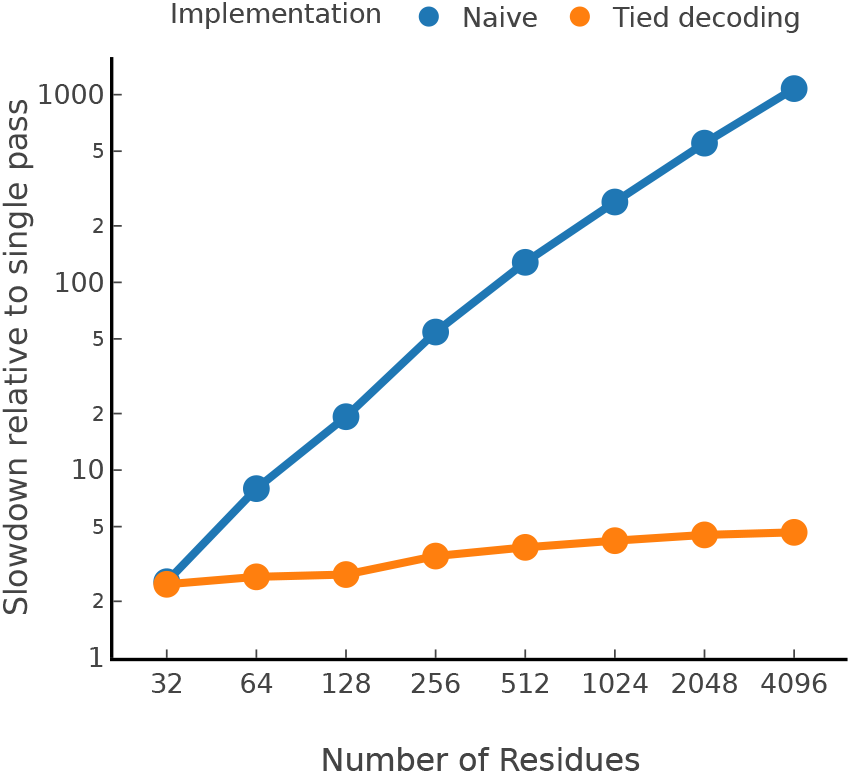
Slowdown relative to a single pass of ProteinMPNN of naïve and tied decoding order schemes

RaSP, DDGun and ACDC-NN have been explicitly trained to reproduce experimental stability data, and they are therefore considered here as supervised models. No experimental protein stability data was used to derive ProteinMPNN and ESMif, and the same applies for ProteinMPNN-ddG and ESMif-ddG which we introduce here. The resulting models are therefore unsupervised.

**Table A.2:**
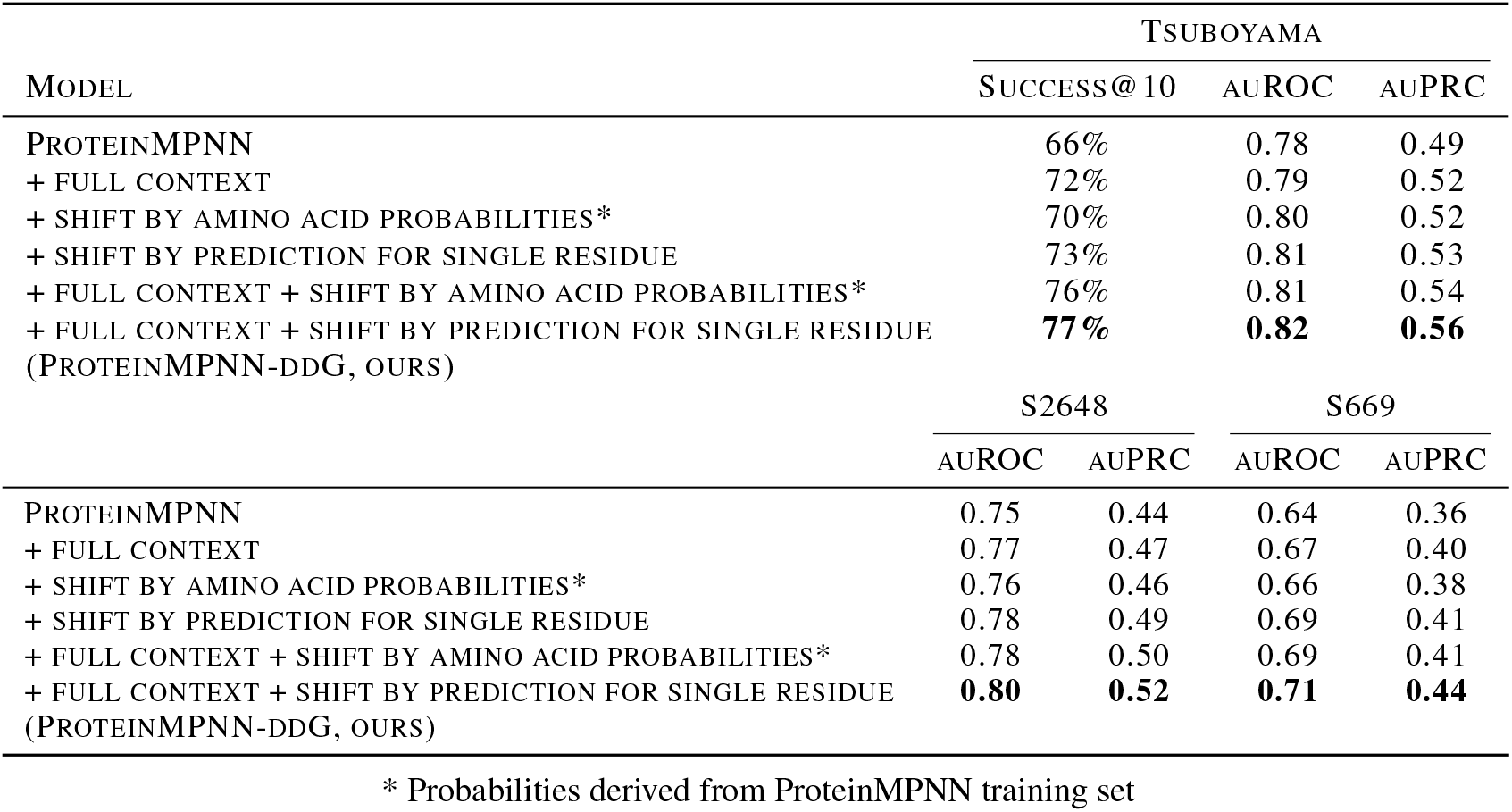
Ablation results for modifications of ProteinMPNN.

## Footnotes

1 https://github.com/compbiomed-unito/acdc-nn, accessed: 16-06-2024

## Notes

https://github.com/PeptoneLtd/proteinmpnn_ddg

### Competing Interest Statement

All authors hold stock options in Peptone Ltd. SB, MI, CF, SR, FA, PF, and KT work for Peptone Ltd.

### Summary of Updates

Small edits and added suggestions to reviewers

